# Agonism between family groups of cooperative breeding smooth-coated otters (*Lutrogale perspicillata*)

**DOI:** 10.1101/2023.12.18.572272

**Authors:** Fan Zheng, Darren Ang Chuin How, Mei-Mei Tan Heng Yee, Philip Johns

## Abstract

Intraspecific agonistic interactions are common in the animal kingdom and have significant consequences for animals. With a growing population and limited resources, these interactions have been observed more frequently among smooth-coated otters in Singapore. However, little is known about the behavioral patterns of these interactions and the factors that may affect their outcomes. To address this, we analyzed intraspecific agonistic interactions through publicly available videos from citizen scientists. Our analysis revealed different behavioral patterns and typical behaviors between winning and losing family groups in different environments. We also found suggestive evidence that larger family groups with more individuals than opponents have an advantage in these interactions. The environment in which an interaction occurs may also affect its intensity.

## Introduction

Agonistic interactions are prevalent in many species of animals (Bell & Currie, 2010; Drummond, 2001; Gobush & Wasser, 2009; Krieg & Burnett, 2017; Leeds et al., 2021, 2022; Scott et al., 2023; Su & Birky, 2007; Wheeler et al., 2013), and usually occur over access to resource such as food, space, and mates (Drummond, 2001; Krieg & Burnett, 2017; Su & Birky, 2007). Agonistic interactions can affect population size, reproductive success, and the formation of dominance hierarchies or territoriality among competitors (Holekamp & Strauss, 2016). Some cooperatively breeding species engage in group territorial defense (Fitzpatrick & Bowman, 2016; Mares et al., 2012; Ridley, 2016). For example, wolves attack other wolves near their dens during the denning season (Smith et al., 2015), which may motivate group territorial defense, and relative pack size increases the chance one pack will displace another (Cassidy et al., 2015). The interplay between resource defense, delayed dispersal and reproductive suppression, kinship and indirect benefits, resource inheritance, and direct benefits can contribute to the evolution of this breeding system (García-Ruiz et al., 2022).

Of 13 extant species of otters (*Lutrinae*), several are social (Lodé et al., 2021), and at least three are cooperative breeders: giant (*Pteronura brasiliensis*), small-clawed (*Aonyx cinereus*), and smooth-coated (*Lutrogale perspicillata*) otters (Bungum et al., 2021; Lélias et al., 2021). Smooth-coated otters typically live in family groups (“romps”) that consist of a breeding pair and their offspring of different ages (Bungum et al., 2021). These groups occupy and defend home ranges that vary in size and quality depending on the availability of food and shelter resources (Estes, 1989; Kruuk, 2006). As a result, agonistic interactions occur sporadically when family groups encounter each other, which can lead to chases, fights, and injuries among the competing otters, especially pups (personal observations, Kruuk, 2006).

In Singapore, smooth-coated otters re-established themselves in 1998 (Khoo & Lee, 2020) after being locally extinct since the 1960s due to the deterioration of the local aquatic ecosystems (Sivasothi & Nor, 1994; Tortajada & Joshi, 2014). There are now approximately 17 family groups and 170 individual smooth-coated otters (Shivram et al., 2023) living in the urban matrix in Singapore. The dense otter population, limited resources, and high degree of urbanization (Shivram et al., 2023; Theng & Sivasothi, 2016) may contribute to competition among family groups. Agonistic interactions have been recorded by otter watchers and citizen scientists on land or in Singapore’s waterways, especially in areas where otter groups have overlapping or adjacent home ranges (Shivram et al., 2023).

Although not uncommon, little is known about agonistic interactions among smooth-coated otters. Here, we provide a preliminary description of behavioral patterns, the differences between winners and losers of agonistic interactions, and factors that may influence outcomes to elucidate an important component of cooperative breeding systems. Video recordings were collected from otter watchers and citizen scientists in Singapore. Citizen science can be effective in animal behavior studies, and some studies glean data from public sources (Boydston et al., n.d.; Bungum et al., 2022; Bungum & Johns, 2022; Loong et al., n.d.). Gleaning data from public repositories can be particularly useful when the behavior is rare or difficult to find (Nelson & Fijn, 2013), like group territorial defense.

## Methods

Intraspecific agonistic interactions are episodic and sporadic among smooth-coated otters, and by focusing on behaviors within these interactions, we hoped to lessen bias from *ad-lib* sampling (Altmann, 1974). We analyzed videos gathered from otter watchers in Singapore and from online sources, including YouTube (www.youtube.com) and Facebook (www.facebook.com) (Table A1). In this study, we excluded: 1) video recordings where both families did not appear together; 2) redundant video recordings where the same interaction was captured from different angles; 3) incomplete video recordings where the winner or loser was unclear; and 4) interactions with fewer than two otters from each family group involved. Thus, we selected 13 video recordings of 12 interactions among different family groups in Singapore (Table A1). Hereinafter, every term “**interaction(s)**” we used in this paper refers to these selected 12 interactions only.

Ethograms from 20 published articles (Table A2) on different otter species were collected to build an ethogram pool (Table A3), from which we selected appropriate behaviors. We added behaviors to our ethogram as we observed them. Agonistic interactions could occur on land, in water, or partly on land and partly in water. Because behavioral patterns of the interactions that happened fully or partly in water were similar, we categorized “**land interactions**” as interactions that occurred entirely on land, while all other types of interactions were categorized as “**water interactions**”. Then, we built ethograms for water (Table A4) and land (Table A5) interactions separately. We also categorized each behavior item as a “**point**” or “**state**” behavior based on whether a behavior is instantaneous or continuous.

We used the program Behavioral Observation Research Interactive Software (BORIS, Friard & Gamba, 2016; http://www.boris.unito.it) to describe behavioral sequences of aggressive encounters, employing water and land ethograms (see Tables A4 and A5). We also made assignations according to the following rules: 1) Group-level behaviors: When more than 70 percent of individuals in a family group perform the same behavior, we record the uniform behavior only once. 2) Winning and losing: The losing family group flees without re-engaging at the end of an interaction; the winning family group chases or follows. Notice that within the interaction, some behaviors involved both sides, which could not be attributed to either family because of the melee and were defined as “**intense**” behaviors (See Tables A4 and A5). 3) The start of an interaction was when the relative distance between two family groups narrowed. The endpoint was the losing family group moving away and not re-engaging.

We exported the integrated behavioral strings from BORIS to the accompanying program BEHATRIX (http://www.boris.unito.it/pages/behatrix) to determine which behavioral transitions occurred more frequently than chance, based on 10,000 permutations and more frequently than one percent of the time. The processed behavioral strings were exported to GraphViz (Ellson et al., 2002; https://graphviz.org/) to generate kinetic flow diagrams for interactions on land and in water.

We counted the number of times each behavior occurred (n) and the percentage of the total length of interaction each state behavior accounted for (% of the total length, or “**duration”**). Interactions varied in length, and to eliminate the effects of this variability, we calculated the “**frequency**” (n/total length of an interaction). State behaviors (Tables A4 and A5) can occur for a long time but may not occur often, and duration accounts for that variability.

We compared the behaviors of the winning and losing family groups in aggregate with a PERmutated Multiple ANalysis Of VAriance (PERMANOVA), using the function *adnois2* (Anderson, 2017; R vegan package, https://github.com/vegandevs/vegan) and 1000 permutations. To account for the pairwise nature of interactions, we nested the behavioral frequencies and durations of winning and losing family groups within individual encounters. In this study, we used a distance matrix (dismatrix) because the research question focused on pairwise dissimilarities (Anderson, 2017). We also conducted a Principal Coordinates Analysis (PCoA) using the function *cmdscale* (Cailliez, 1983; R core team, https://www.R-project.org) to visualize differences in behavioral patterns between the winning and losing family groups.

We compared the frequencies and durations of individual behaviors between winning and losing family groups within each encounter. To identify significant differences, we calculated pairwise mean differences between winning and losing family groups in both frequency and duration for all behaviors, along with the bootstrapped 95% confidence intervals, using the function *boot* (Davison & Hinkley, 1997; https://CRAN.R-project.org/package=boot). We highlighted behavioral flow diagrams with behaviors performed with higher frequency or duration by the winning and losing family groups, identified through pairwise differences.

In some encounters, the losing family group did not engage with the winning family group, either running away immediately upon seeing the other family group or being overtaken by the winning family group without showing aggressive behaviors (i.e., **non-aggression**). In other encounters, the losing family group actively and aggressively engaged with the winning family group (i.e., **aggression**).

To explore the potential factors affecting the aggressiveness of the losing family groups, we considered the demographic factors and the environment in which the interactions took place via generalized linear models (GLMs) using function *glm* (Cailliez, 1983, R core team, https://www.R-project.org). In this descriptive study, we introduced various models and highlighted the similarities among them (Burnham et al., 2011). After this, the inflection points of the prediction curves of the selected models were calculated using function *ese* (Christopoulos, 2012, inflection package, https://cran.r-project.org/web/packages/inflection/vignettes/inflection.html).

## Results and Discussion

### Behavioral patterns

PERMANOVA revealed aggregate pairwise differences in behavior frequencies and durations between winning and losing otter families in water interactions but not land interactions (Table 1). PCoA plots also demonstrate that the overall behavioral differences between winning and losing family groups were clearer in water interactions (Figure 1. 1a, 1b) than land interactions (Figure 1. 2a, 2b).

**Table 1.** PERMANOVA results for the behaviors among interactions by the winning and losing family groups.

| Item | Environment | Source | df | SS | MS | Pseudo-F | R<sup>2</sup> | p-value |
| --- | --- | --- | --- | --- | --- | --- | --- | --- |
| Frequency | Water | Family (W/L) | 1 | 0.024 | 0.024 | 5.589 | 0.285 | **0.008 \*\*** |
|  |  | Residuals | 14 | 0.059 | 0.004 |  |  |  |
|  |  | Total | 15 | 0.083 |  |  |  |  |
| Duration | Water | Family (W/L) | 1 | 4735.3 | 4735.3 | 6.118 | 0.304 | **0.008 \*\*** |
|  |  | Residuals | 14 | 10835.1 | 773.9 |  |  |  |
|  |  | Total | 15 | 15570.4 |  |  |  |  |
| Frequency | Land | Family (W/L) | 1 | 0.002 | 0.002 | 1.608 | 0.211 | 0.250 |
|  |  | Residuals | 6 | 0.007 | 0.001 |  |  |  |
|  |  | Total | 7 | 0.009 |  |  |  |  |
| Duration | Land | Family (W/L) | 1 | 1493.7 | 1493.7 | 1.042 | 0.148 | 0.375 |
|  |  | Residuals | 6 | 8603.6 | 1433.9 |  |  |  |
|  |  | Total | 7 | 10097.2 |  |  |  |  |

**Figure 1.**
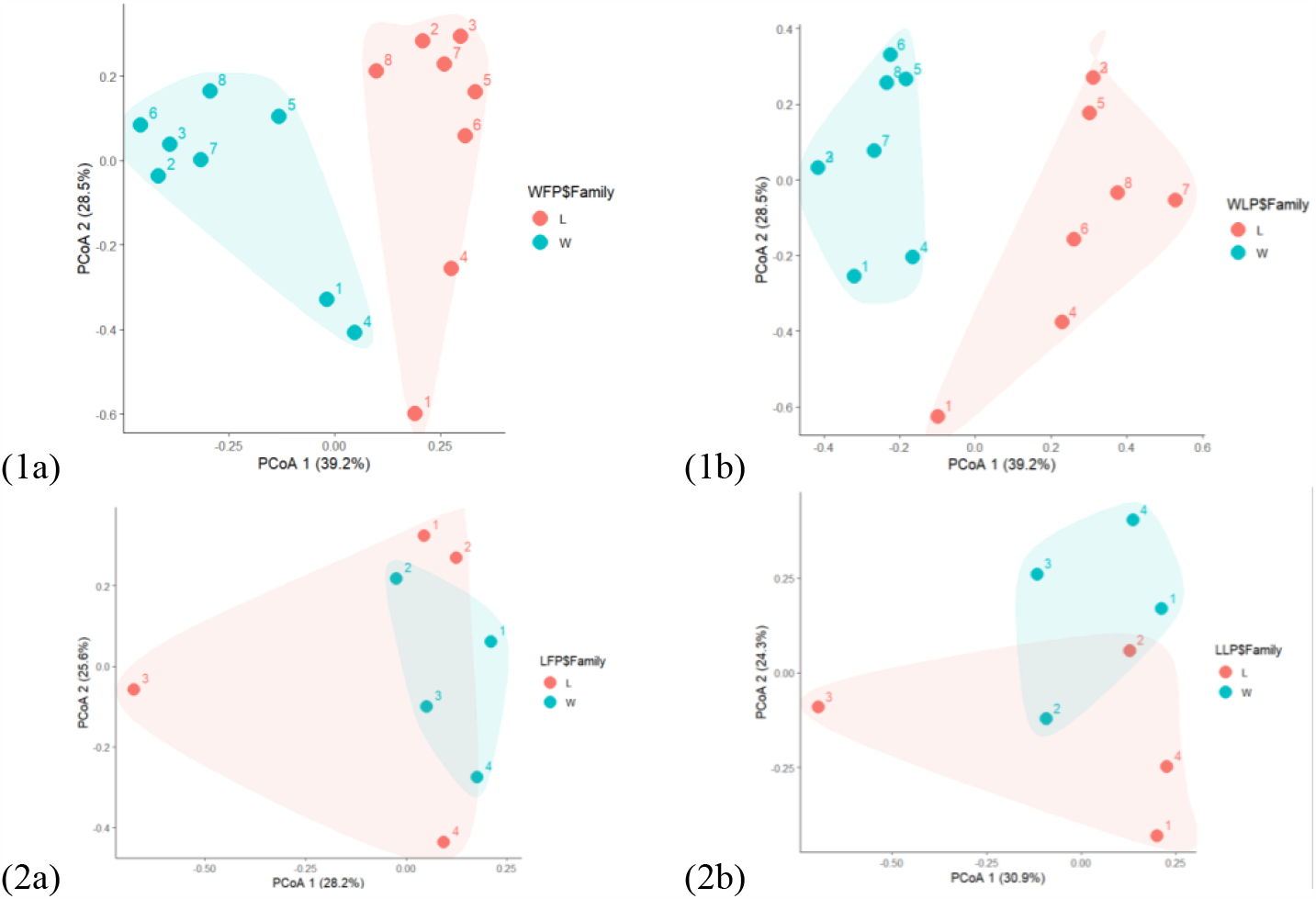
PCoA plots of behavior frequencies and durations in winning and losing family groups during water and land interactions. Land frequency (1a) and duration (1b), and water frequency (2a) and duration (2b) of behaviors. Numbers identify individual interactions for comparing winners (light green) and losers (light orange; see Table A2.) X and Y are principal coordinate axes with percentage values indicating differences in sample composition.

**Figure 2.**
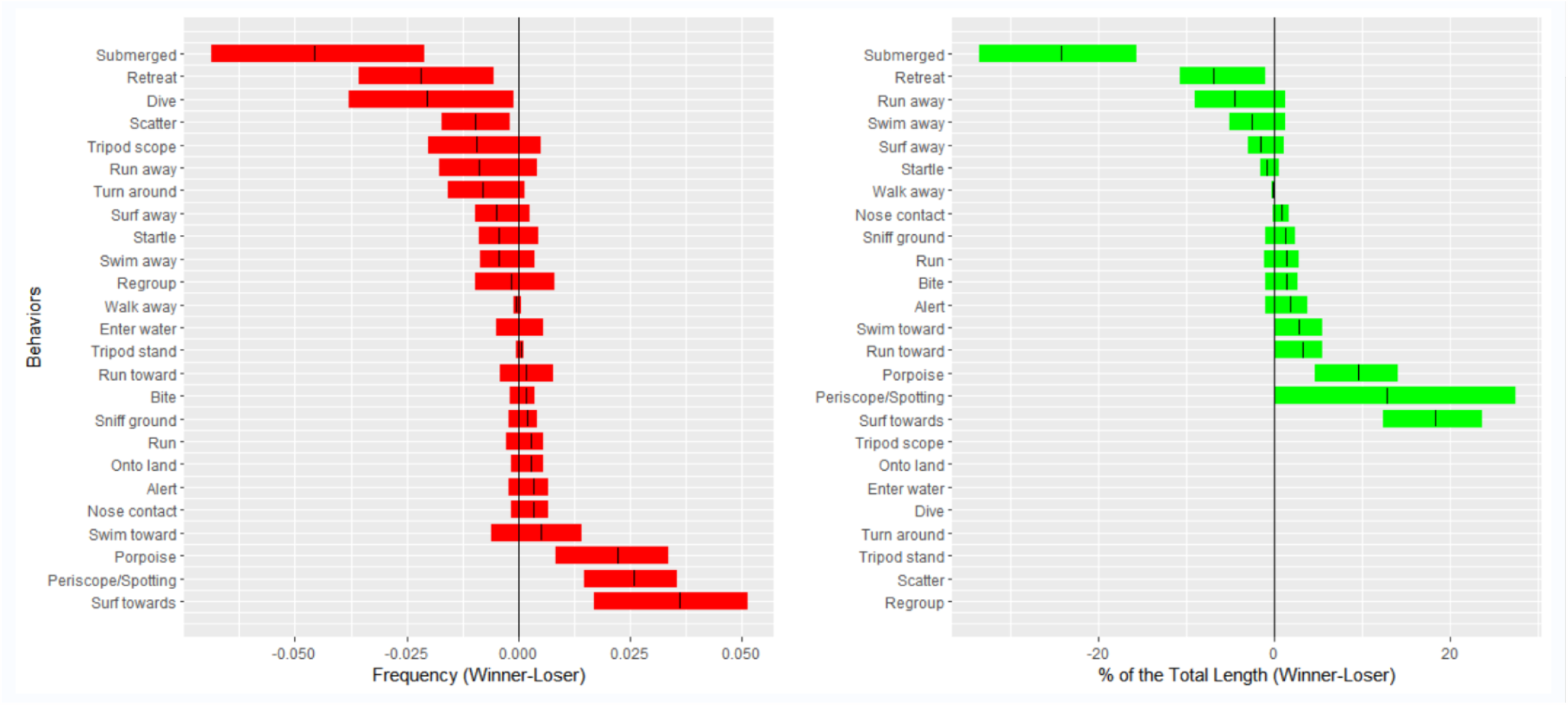
Pairwise differences in behavioral frequencies and durations between winning and losing family groups in water. Means (lines) and 95% confidence intervals (bars) for frequencies (red) and durations (green) of behaviors. Means > 0 indicated typical behaviors for winning family groups; < 0 for losing family groups. Point behaviors could not be included in durations (green).

Pairwise differences for water interactions between winning and losing family groups, within agonistic interactions, in frequency and duration of each behavior, revealed typical differences (Figure 2). Losing family groups tended to submerge, retreat, dive, and scatter more frequently than winners while winning family groups tended to porpoise, periscope/spot, and surf toward the losers more frequently. Losing family groups spent a greater percentage of time submerged or retreating while winning family groups spent a greater percentage of time porpoising, periscope/spotting, or surfing towards the losers. Although PERMANOVA revealed no significant difference between winning and losing family groups on land (Table 1), we still noted typical differences among land interactions (See Figure A1.).

Behavioral flow diagrams of interactions in water (Figure 3) and on land (Figure A2), with highlighted typical and intense behaviors, revealed common behavioral transitions. Notice “loops” among behavioral transitions’ common patterns and all typical behaviors of winning and losing family groups were included in some loops, emphasizing transitions among them that occur more frequently than chance. The winning family group employed searching behaviors like periscope spotting, then swam very quickly at the other family with surfing and porpoising behaviors. The losing family group responded by retreating, diving, and submerging.

**Figure 3.**
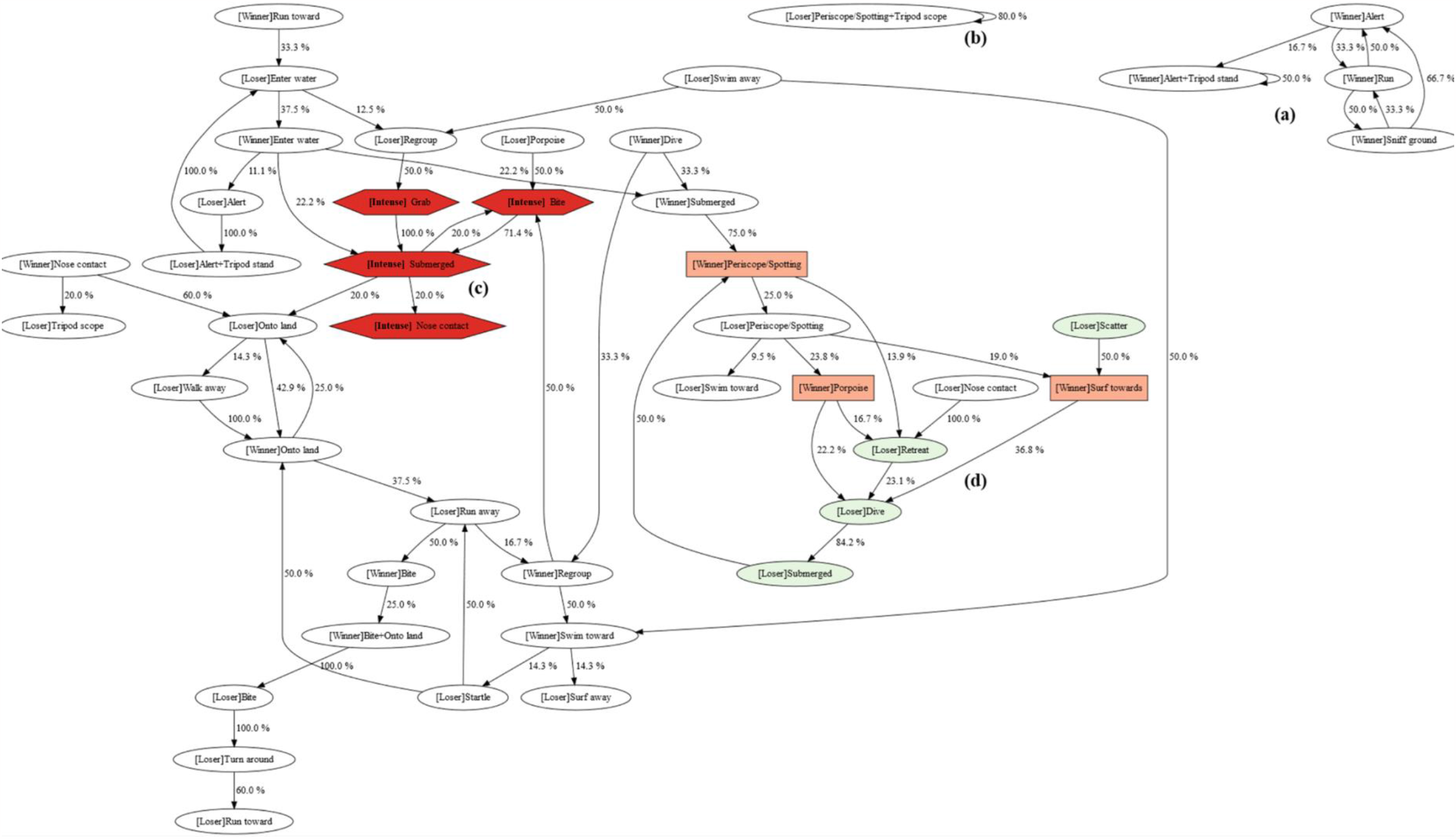
Behavioral sequences of water interactions. Arrows indicate the proportion of observed transitions. The family group (Winner, Loser) of the actor is in brackets for ordinary behaviors (ovals); in intense behaviors (red hexagons), the family origin of actors was unclear; + indicates simultaneous behaviors. Other colors indicate typical behaviors of the winning (light orange) and losing (light green) family groups (Figure 2; see Table A4 for ethogram).

In interactions where we had video of the whole process, one otter family appeared to search for the opponent early in the interactions through visual (e.g., alert, tripod stand; see Figure 3 and Figure A2, (a), (b)) and scent (e.g., sniff ground; see Figure 3 and Figure A2, (a), (b)). The winning family was generally more active in searching and employed more searching behaviors.

In some encounters (6 of 12), otter families interacted directly with extensive physical contact, such as wrestling, grabbing, and biting (Figure 3 (c); see Fig A2 (c)). These looping cycles of behaviors could occur repeatedly. On land, one family often attacked the other, but in water, family groups typically swam towards each other initially. Losing families tended to scatter after the physical interactions but sometimes regrouped or fought back in subgroups.

For both in water and on land, otters touched noses after direct physical contact (Figure 3(c); see Fig A2 (c)). Violent physical fights could cause otter family members to mix, and otters seemed to use nose contact to gauge which family another otter belonged to. After fierce interactions, both winning and losing families would touch noses. Losing family groups performed nose contact before quickly and fleeing the winning family.

In water interactions, winning family groups visually searched for opponents, while losing family groups monitored the opponent’s movements (periscope/spotting, Figure 2 (d); see Table A4). After locking onto the opponent’s position, the winning family group “surf towards/porpoise” where the losing family groups were. However, the losing family groups tended to “retreat” or “submerged” to avoid contact by making them invisible and no longer in their original position. So, the winning family groups lost their target and did “Periscope/spotting” to try to lock onto the losing family again. When the losing family groups emerged from the water, the winning family groups could lock onto their targets again and repeat the sequence of behaviors. For land interactions, we also observed looping cycles of behaviors where winning families chased opponents several times before losing families finally ran away (e.g., Figure A2, (d)). However, the behavioral patterns were much less complicated than those of water interactions. This may be because aquatic environments posed more opportunities for otters to display their dominance or submission, e.g., by diving to get away. It should be noted that limited data existed for the land interactions, which could also be a reason (3 of the 12 interactions). The observed behavior looping cycles were highly structured, providing insights into animal communication systems, social dynamics, and environmental adaptation strategies, which can inform conservation efforts.

Larger otter family groups won 92% (11 of 12) of recorded interactions (X^2^=8.33, p = 0.0039). We performed GLMs to determine what influenced whether the losing family would engage in an aggressive interaction, and we included as main effects the difference in winner and loser family headcounts, the number of individuals in the losing family, the ratio winning to losing family headcounts, and the environment (Table A6). The best model (Model 1; ∂BIC 3.30) included only the number of individuals in the losing family as a significant predictor of aggression (Table 2).

**Table 2.** Generalized linear models 1–6 predict the aggressive level of the family groups that finally lost the interaction against the opponent. Coefficient estimates, standard errors, z-values, and p-values for each predictor; AIC and BIC for each model. See Table 9 for the description of variables.

| Model | Model 1 |  |  |  | Model 2 |  |  |  | Model 3 |  |  |  |
| --- | --- | --- | --- | --- | --- | --- | --- | --- | --- | --- | --- | --- |
| Predictors | Estimate | Std. Error | z value | Pr (>\|z\|) | Estimate | Std. Error | z value | Pr (>\|z\|) | Estimate | Std. Error | z value | Pr (>\|z\|) |
| (Intercept) | 4.109 | 2.361 | 1.741 | 0.082 | -1.996 | 1.299 | -1.537 | 0.124 | -1.996 | 1.454 | -1.373 | 0.170 |
| spe |  |  |  |  | 2.030 | 1.230 | 1.651 | 0.099 |  |  |  |  |
| L | -0.880 | 0.503 | -1.748 | 0.080 |  |  |  |  |  |  |  |  |
| dev |  |  |  |  |  |  |  |  | -0.536 | 0.370 | -1.447 | 0.148 |
| envWF |  |  |  |  |  |  |  |  |  |  |  |  |
| AIC |  | **14.851** |  |  |  | 15.101 |  |  |  | 16.839 |  |  |
| BIC |  | **15.820** |  |  |  | 16.071 |  |  |  | 17.809 |  |  |
| ∂BIC |  | **3.300** |  |  |  | 3.049 |  |  |  | 1.311 |  |  |

| Model | Model 4 |  |  |  | Model 5 |  |  |  | Null Model |  |  |  |
| --- | --- | --- | --- | --- | --- | --- | --- | --- | --- | --- | --- | --- |
| Predictors | Estimate | Std. Error | z value | Pr (>\|z\|) | Estimate | Std. Error | z value | Pr (>\|z\|) | Estimate | Std. Error | z value | Pr (>\|z\|) |
| (Intercept) | -2.461 | 1.794 | -1.372 | 0.170 | 1.067 | 3.547 | 0.301 | 0.764 | 0.000 | 0.577 | 0 | 1 |
| spe | 2.059 | 1.275 | 1.615 | 0.106 |  |  |  |  |  |  |  |  |
| L |  |  |  |  | -0.609 | 0.492 | -1.238 | 0.216 |  |  |  |  |
| dev |  |  |  |  | -0.324 | 0.421 | -0.770 | 0.441 |  |  |  |  |
| envWF | 0.648 | 1.622 | 0.399 | 0.690 | 0.913 | 1.727 | 0.529 | 0.597 |  |  |  |  |
| AIC |  | 16.937 |  |  |  | 17.835 |  |  |  | 18.636 |  |  |
| BIC |  | 18.392 |  |  |  | 19.774 |  |  |  | 19.120 |  |  |
| ∂BIC |  | 0.728 |  |  |  | -0.654 |  |  |  | - |  |  |

Larger families are more likely to engage in aggressive interactions. The inflection point for the best model was 4.5 (see Figure A3. a), suggesting that families with fewer than five members tend not to escalate and rather run away. The second-best model (Table 2, Model 2; ∂BIC = 3.05) provided weak evidence that, as the headcount difference between the winning and losing family groups decreased, the smaller family group tended to engage aggressively. That inflection point was -3.3 (Figure A3.b), suggesting that when a larger family group had more than three additional individuals than a smaller family group, the smaller family group was likely to give up without engaging the larger family directly; they thus became the losing family group. How otters assessed the size of their family groups relative to an opponent’s, i.e., through visual cues, auditory cues, or long-term exposure to sprain and olfactory cues, was beyond the scope of this study.

Our findings suggested that a young breeding pair (two adults) with only one brood of offspring (about three pups) is probably not big enough to defend their territory successfully. Similar to wolves (Smith, 2015), pup mortality is sometimes a consequence of smooth-coated otter agonistic interactions (Khoo & Sivasothi, 2018). Agonistic interactions, while sporadic, are not rare. Although smooth-coated otters engage in several group behaviors, including group defense against predators (Bungum & Johns, 2022), defense of group territories against conspecifics may be what drives the delayed dispersal of older broods in this cooperatively breeding otter.

Agonistic interactions are influenced by factors other than demographics. Aggressiveness among different individuals varies; some individuals can deviate from the behaviors of the group. Factors such as hormone levels, age, sex, body size, status within the group, and early-rearing conditions may also influence an individual’s aggressive behavior (Holekamp & Strauss, 2016; Mikkola et al., 2021). At the group level, the family structure may influence aggressiveness. The presence of pups could increase the aggression of a group (Bungum & Johns, 2022; Dyble et al., 2019), or it may lead to families being more defensive and less aggressive. However, our limited data did not support further discussion about these factors. Future studies could focus on clarifying the influence of population factors and studying more potential factors, especially the social structure of otter family groups and the differences among water and land interactions.

## Acknowledgements

We would like to thank the otter-watching community of Singapore, especially Chun Kit Soo, Kennie Goh, Muhammad Harith, Otter Kwek, and participants on the Facebook groups Otter Channel and Otter City for their help with this study. Portions of this study were conducted by Fan in partial completion of MSC requirements for BCNBCS, NUS; portions were conducted by Tan in partial completion of Capstone requirements, and by Ang in partial completion of Summer Research requirements at Yale-NUS College. The work was supported in part by the Singapore Ministry of Education through the Yale-NUS College grant R-607-265-226-121, by Yale-NUS College Centre for International and Professional Experience (CIPE), and by Singapore NParks permits NP/RP20-073 and NP/RP20-075.

## Notes

**Conflicts of interest:** The authors have no competing financial or non-financial interests that are directly or indirectly related to this study.

https://doi.org/10.6084/m9.figshare.24848316.v3

### Competing Interest Statement

The authors have declared no competing interest.

