## Supplementary material for "Agonism between family groups of cooperative breeding smooth-coated otters (*Lutrogale perspicillata*)": Figures A1 to A3

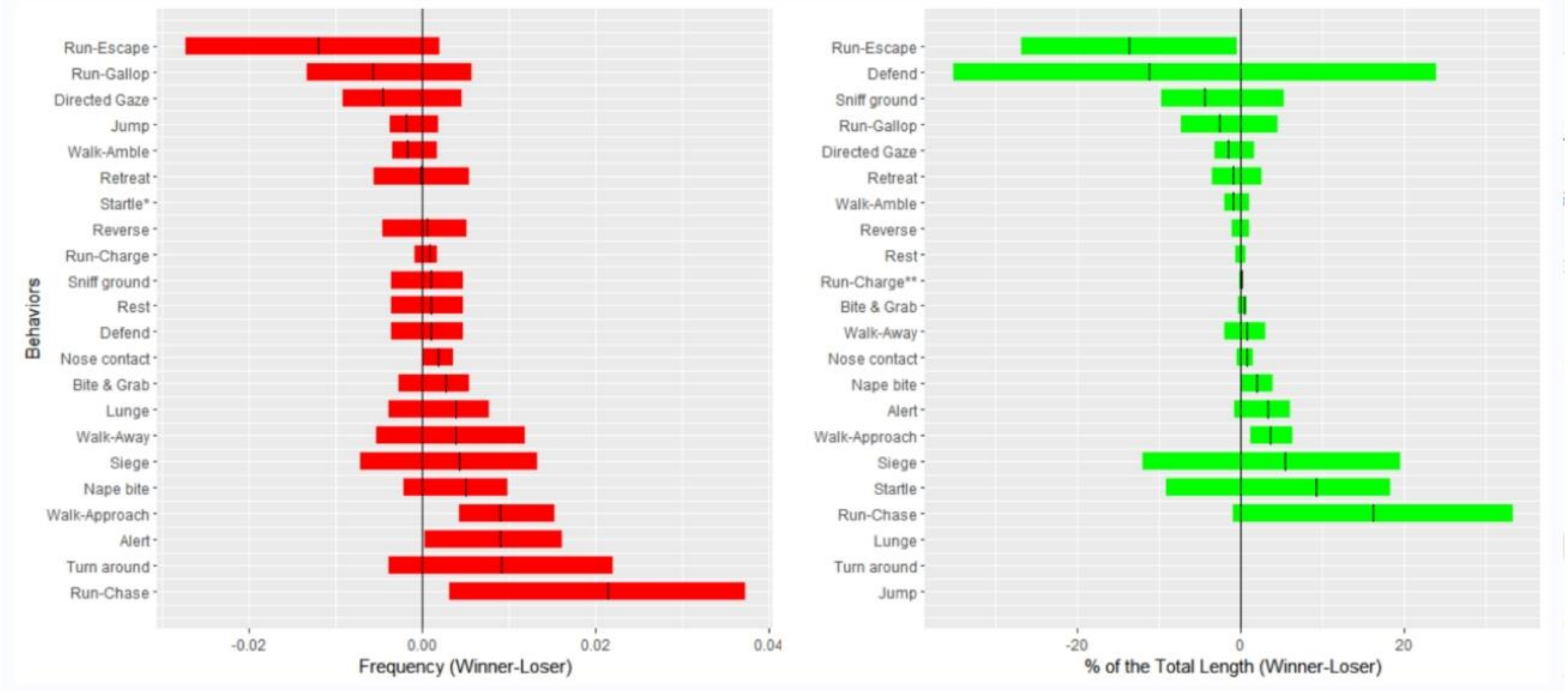


Figure A1. **Pairwise differences in behavioral frequencies and durations between winning and losing family groups on land.** *Means (lines) and 95% confidence intervals (bars) for frequencies (red) and durations (green) of behaviors. Means > 0 indicated typical behaviors for winning family groups; < 0 for losing family groups. Point behaviors could not be included in durations (green).*

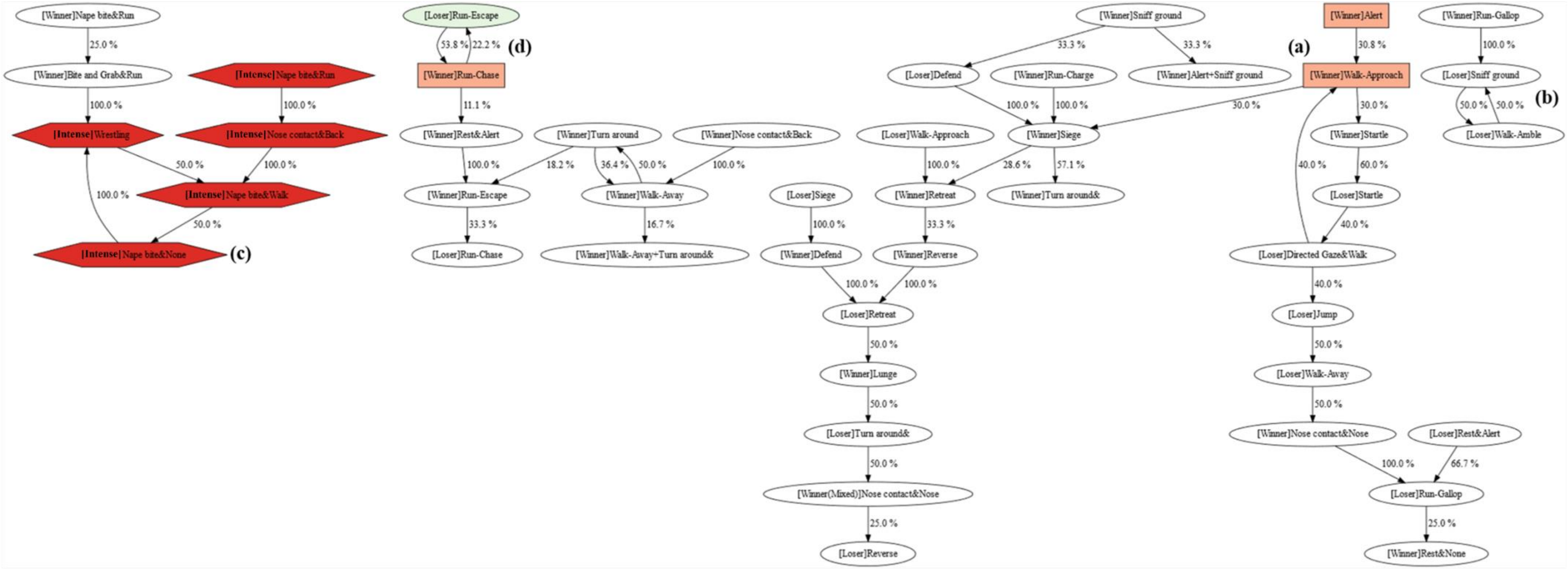


Figure A2. **Behavioral sequences of land interactions.** *Arrows indicate the proportion of observed transitions. The family group (Winner, Loser) of the actor is in brackets for ordinary behaviors (ovals); in intense behaviors (red hexagons), the family origin of actors was unclear; + indicates simultaneous behaviors. Other colors indicate typical behaviors of the winning (light orange) and losing (light green) family group (Figure A1; see Table A5 for ethogram).*

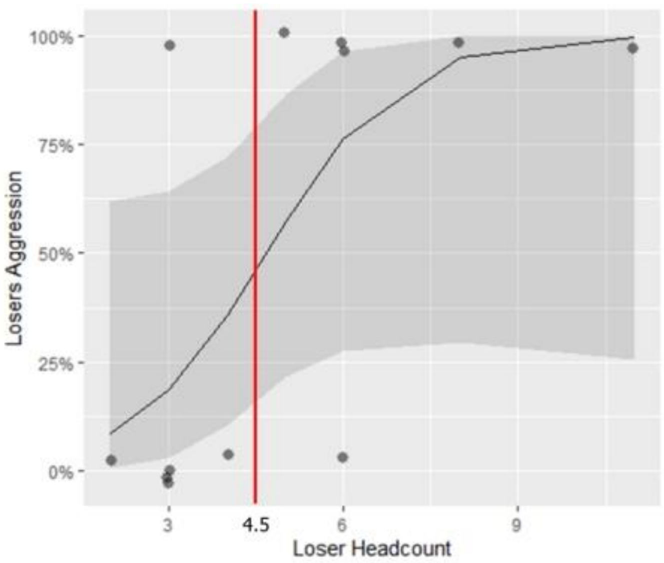


(a) The best-fitting model: Model 1

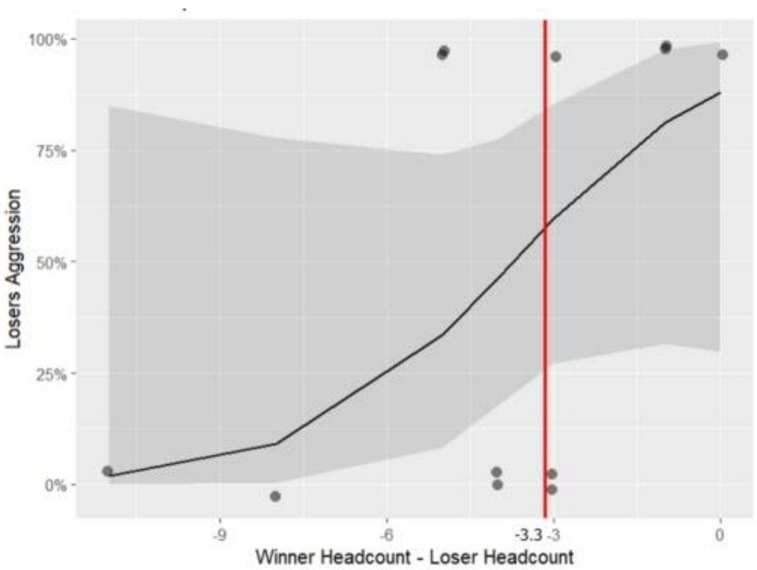


(b) The better-fitting model: Model 2

Figure A3. **Predicting losing family groups' aggression in response to ppponents in the interactions.** *Panels a, b, c - Models 1, 2, 6. Y-axis – aggression of losing family groups. Panel a: X-axis - losing family groups' headcount. Panels b, c: X-axis - headcount difference between winning and losing family groups. Panels a, b - land and water interactions combined. Panel c: land (dotted lines) and water (solid lines) interactions separate. Gray area - 95% confidence intervals. Red line - inflection point of prediction curve, x coordinates marked on x-axis.*
