## Supplementary material for "Agonism between family groups of cooperative breeding smooth-coated otters (*Lutrogale perspicillata*)": Table A5

Table A5. Ethogram describing behaviors scored in otters' intraspec
multiple otter species from previous studies (See Table A2 for the sc
this ethogram is completely new. The behaviors were categorized in

| Type[+] | Key | Code | Category |
|---|---|---|---|
| Point behavior/event | a | Bite (fight) | Land |
| State behavior/event | b | Bite & Grab* | Land |
| State behavior/event | c | Nape bite** | Intense |
| State behavior/event | d | Wrestling | Intense |
| State behavior/event | e | Alert | Land |
| State behavior/event | f | Nose contact*** | Intense |
| State behavior/event | g | Startle | Land |
| State behavior/event | h | Directed Gaze**** | Land |
| Point behavior/event | i | Turn around | Land |
| State behavior/event | j | Run-Gallop | Land |
| State behavior/event | k | Run-Escape | Land |
| State behavior/event | l | Run-Chase | Land |
| State behavior/event | m | Run-Charge | Land |
| State behavior/event | n | Walk-Amble | Land |
| State behavior/event | o | Walk-Away | Land |
| State behavior/event | p | Walk-Approach | Land |
| Point behavior/event | q | Flinch away | Land |
| Point behavior/event | r | Lunge | Land |
| State behavior/event | s | Defend | Land |
| State behavior/event | t | Siege | Land |
| State behavior/event | u | Rest | Land |
| Point behavior/event | v | Reverse/Back-up | Land |
| State behavior/event | w | Retreat | Land |
| State behavior/event | x | Sniff ground | Land |

[illegible]
* Modifiers of behavior item "Bite & Grab": Run, Walk; based on tl
** Modifiers of behavior item "Nape bite": Run, Walk; based on the
*** Modifiers of behavior item "Nose contact": Nose, Back; based (

[illegible] Modifiers of behavior item “Nose contact”: Nose, Back; based [illegible]

**** Modifiers of behavior item “Directed Gaze”: Run, Walk; base[illegible]

+ The term "event" is the default for BORIS. Since in the article we [illegible] ific agonistic interactions on land. This ethogram builds solely on the collected ethograms for ources of the studies, and see Appendix for the collected ethograms). Unlike water ethogram, to: Interaction and Land.

**Description**

Otter opens mouth and closes on another otter intensely and briefly
Otter closes mouth on the body of another another, maintaining prolonged contact.
Otter closes mouth around nape of neck of another otter.
Otters actively grasp each other with forearms around the head and shoulders, then twist to break the hold
Otter stop mid-activity, raising head to look around, sometimes standing in an upright position on their hind legs to survey the area.
Otter uses nose to touch another otter in a solicitous, usually the nose but can be on any part of the body.
Otter pauses mid activity and makes direct eye contact with stimuli
Otter makes direct eye contact with stimuli.
Otter turns ~180 degrees to face the opposite direction
Otter moves at fast speed, slightly jumping, alternating anterior with posterior limbs
Otter moves at fast speed, slightly jumping, alternating anterior with posterior limbs away from object/animal.
Otter moves at fast speed, slightly jumping, alternating anterior with posterior limbs towards fleeing object/animal.
Otter moves at fast speed, slightly jumping, alternating anterior with posterior limbs towards stationary object/animal.
Otter moves at slow speed.
Otter moves at slow speed away from object/animal.
Otter moves at slow speed towards object/animal, may stop periodically during approach, and stopping once near.
Otter swerves or jerks its body away from another immediate otter, sometimes taking a few steps back.
Otter suddenly performs a forward body movement towards another immediate otter
Otter positions self to block entrance to holt from other otters, vocalizing and posturing, while remaining within holt.
Otter(s) surround other otter(s) in holt, vocalizing and posturing without attempting to enter holt.
Otter is stationary with paws and body on ground, head raised above ground.
Otter steps backwards with head/neck pushed down but nose tilted up, oriented towards conspecific and generally whinning and/or screaming. Can be submissive response to conspecific.
Otter backs away from object/animal while watching it.
Otter lowers nose to ground, moving head back and forth, either while stationary or walking

ne adjoint behaviors
e adjoint behaviors
on the part of the body the action is on

on the part of the body the action is on
d on the adjoint behaviors
classify all the content related to behaviors using the term "behavior(s)", here we also indicate the
